# Impact of GPR39 Inhibition on Tissue Oxygen Tension in Acute Myocardial Infarction: Modeling Using an Indirect Response Pharmacokinetics/Pharmacodynamics Approach

**DOI:** 10.64898/2026.09.24.754288

**Authors:** Davide Ronchi, Chiara Roversi, Simone Zannoni, Alessandro De Carlo, Roberto Visentin, Annalisa Pellacani, Lisa Morbioli, Jessica Franchi, Carmen Methner, Marco Pergher, Alessia Tagliavini, Alberto Parazzoli, Sanjiv Kaul

## Abstract

In pre-clinal studies, VC108, a novel GPR39 inhibitor, increased myocardial O_2_ tension (mpO_2_) during different phases of acute myocardial infarction (AMI). The purpose of this work was to investigate the temporal effects of VC108 on mpO_2_ using a novel population PK/PD (popPK/PD) model developed from pre-clinical data using healthy animals and animals subjected to AMI protocols. PK/PD simulations were then performed to explore mpO_2_ dynamic in untested experimental scenarios. VC108 PK data fitted well to a two-compartment model with linear elimination with body weight, sex, and nominal dose identified as significant covariates influencing clearance and/or central compartment volume. Applying indirect effect model principles for representing mpO_2_ flux, a structural popPK/PD model was developed to capture mpO_2_ dynamics during different AMI phases: before and during coronary occlusion and after reperfusion. The model successfully captured basal mpO_2_ homeostasis, ischemic decline, and reperfusion overshoot, while quantifying the contribution of VC108 to collateral blood flow and functional recovery. Dose-response relationships were simulated to measure the effect of different VC108 dose levels on permanent damage, i.e. a reduction in mpO_2_ at steady state after AMI. The probability of remaining below predefined damage thresholds (mpO_2_ reduction of 5% to 20%) increased with dose in both sexes, with females consistently achieving greater protection that plateaued around 1□mg/kg of VC108 for higher permanent damage. This model represents a valuable tool to describe, for the first time, VC108 efficacy data and pharmacokinetics with ischemia–reperfusion dynamics and quantify its effect in untested dose levels.

## Introduction

Acute myocardial infarction (AMI) is one of the leading causes of death and disability worldwide [1]. Although timely reperfusion therapies have significantly improved outcomes [2], many patients continue to experience impaired myocardial recovery due to microvascular dysfunction and inadequate O_2_ delivery to ischemic tissue [3]. Restoration of epicardial coronary flow does not always translate into sufficient tissue oxygenation because of persistent reduction in capillary perfusion (no reflow) resulting in larger infarctions with adverse left ventricular (LV) remodeling [4]. Lack of healthy tissue within the risk area leads to infarct expansion, ventricular arrhythmias, heart failure, and death [5]. The temporal dynamics of myocardial O_2_ tension (mpO_2_) during ischemia and reperfusion could be used as benchmarks for developing therapeutic strategies for AMI [6].

mpO_2_ within the area at risk for necrosis after acute coronary occlusion is influenced by the magnitude of collateral blood flow (cBF) [7-11]. The spatial extent of cBF at the time of coronary occlusion determines the spatial extent of tissue necrosis measured several hours later [12]. Whereas the entire myocardium within the risk area after coronary occlusion is dysfunctional [13], regions with >20-25% of basal blood flow usually survive necrosis [14]. Hence, any intervention that will deliver >25% of basal blood flow will result in tissue salvage.

When a coronary artery in a normal subject is temporarily occluded for 20 seconds, coronary blood flow increases 4 to 5-fold compared to baseline after release of the occlusion (hyperemic response). Hyperemia is also noted after reperfusion in AMI that lasts for several hours to days [15-18], the magnitude of which is inversely related to myocardial necrosis. mpO2 levels after reperfusion rise and fall in proportion to tissue blood flow [9,10].

Recent studies have identified the orphan G protein-coupled receptor 39 (GPR39) as a key modulator of coronary blood flow [19]. GPR39 is expressed in vascular smooth muscle cells and pericytes, where its activation by the endogenous agonist 15-hydroxy-eicosatetraenoic acid (15-HETE) promotes vasoconstriction through the mobilization of intracellular calcium [20], thus reducing myocardial blood flow. Genetic and pharmacological deletion of this pathway in a mouse AMI model has been shown to alleviate pericyte-mediated constriction, thereby improving microvascular perfusion [21]. Similarly, pharmacological inhibition of GPR39 in a rat AMI model demonstrated that mpO_2_ is favorably affected during coronary occlusion resulting in less no reflow and necrosis after reperfusion [22]. This was achieved using a novel selective GPR39 blocker, VC108, with pharmacokinetic properties suitable for intravenous administration [23].

These findings provided a rationale for investigating GPR39 blockade as a therapeutic strategy to restore mO_2_ delivery during AMI. However, the regulation of mpO_2_ is the result of multiple, time-dependent processes, where drug exposure, microvascular perfusion (both anterograde and through collaterals [24,25] and tissue O_2_ balance interact in a nonlinear manner. To disentangle these factors a quantitative framework is required.

Pharmacokinetic/pharmacodynamic (PK/PD) modelling provides such a framework [26]. By linking drug concentrations to dynamic biological responses, PK/PD models allow the characterization of dose–response relationships as well as the exploration of mechanisms of action and dosing scenarios that have not been directly tested experimentally. In addition to being descriptive, these models are also predictive, supporting simulations of alternative regimens, optimization of therapeutic strategies, and translation from preclinical studies to clinical applications.

PK/PD approaches have already provided valuable insights in cardiovascular medicine. Examples include quantifying the impact of inotropes on cardiac output [27,28], vasodilators on arterial blood pressure [29,30], and inflammatory mediators on reperfusion injury [31,32]. Applying the same workflow to VC108 is particularly relevant since preclinical studies have demonstrated that VC108 improves microvascular perfusion and myocardial O_2_ availability during AMI.

In this work, a population PK/PD (popPK/PD) model was developed based on preclinical data from healthy rats and those subjected to AMI to investigate the impact of VC108 on mpO_2_ and functional recovery, thereby providing a rigorous basis for evaluating its therapeutic benefits. Specifically, starting from indirect effect model principles which represent mpO_2_ flux, a novel popPK/PD model was developed to capture mpO_2_ dynamics during the different phases of AMI: before and during coronary occlusion, as well as after reperfusion. Based on the developed model, PK/PD simulations are then performed to explore mpO_2_ dynamic in untested experimental scenarios to support further evaluations of drug efficacy.

## Material and Methods

### AMI animal study

AMI was induced in Sprague Dawley rats by ligation of the left coronary artery for one hour (coronary occlusion phase), followed by one hour of reperfusion [21]. Animals were randomized to receive either the selective GPR39 antagonist VC108 or vehicle, administered as an intravenous bolus at various time points in relation to coronary occlusion and reperfusion. Hemodynamic parameters, LV wall thickening (echocardiography [33]), and mpO_2_ were continuously assessed. mpO_2_ was quantified using a phosphorescent probe, enabling real-time evaluation of oxygenation in ischemic and non-ischemic regions. These probes have high accuracy and sensitivity allowing measurements as low as 0 Torr [34,35]. Plasma concentrations of VC108 were measured at predefined time points by mass spectrometry.

### PK/PD data summary

The PK/PD model was developed by integrating data from several preclinical studies, where pharmacokinetic and pharmacodynamic were investigated in experimentally induced AMI rats and healthy animals to assess whether drug distribution and response were affected by the disease conditions. A total of 71 Sprague Dawley rats of both sexes were included in 5 different PK studies, with PK sampling comprising an average of about 7 time-points per animal. Data from 4 studies conducted in healthy animals were used, while one AMI study was conducted in rats. Table 1 and Figure 1 provide a comprehensive summary of the origins of the PK data.

**Table 1.** PK data summary. For each study, the specific readout, sampling schedule, tested regimens, animal health status and number per sex are reported.

| Study | Readout | Sampling schedule (hours) | Regimen | Animal health status | Animal number per sex |
| --- | --- | --- | --- | --- | --- |
| 1 | Plasma | 0.083, 0.25, 0.5, 1, 2, 6, 8, 24 | 0.3 mg/kg SD<br>1.0 mg/kg SD<br>3.0 mg/kg SD | Healthy | 3 M<br>3 M<br>3 M |
| 2 | Plasma | 0.083, 0.25, 0.5, 1, 2, 3, 6, 24 | 1.0 mg/kg SD | Healthy | 3 M |
| 3 | Plasma | Day 4, Group 1: 0.083, 1, 3, 24<br>Day 4, Group 2: 0.25, 2, 6 | 3.0 mg/kg QD x 4 days<br>10.0 mg/kg QD x 4 days<br>30.0 mg/kg QD x 4 days<br>40.0 mg/kg QD x 4 days | Healthy | 4M, 4F<br>4M, 4F<br>4M, 4F<br>4M, 4F |
| 4 | Plasma | Day 1: 0.083, 0.25, 0.5, 1, 2, 3, 6, 24<br>Day 14: 0, 0.083, 0.25, 0.5, 1, 2, 3, 6, 24 | 4.0 mg/kg QD x 14 days<br>15.0 mg/kg QD x 14 days<br>40.0 mg/kg QD x 14 days | Healthy | 3M, 3F<br>3M, 3F<br>3M, 3F |
| 5 | Blood | 0.083, 0.25, 0.5, 1 | 0.1 mg/kg SD<br>0.3 mg/kg SD<br>1.0 mg/kg SD | Acute Myocardial Infarction | 3 M<br>3 M<br>3 M |

**Fig. 1.**
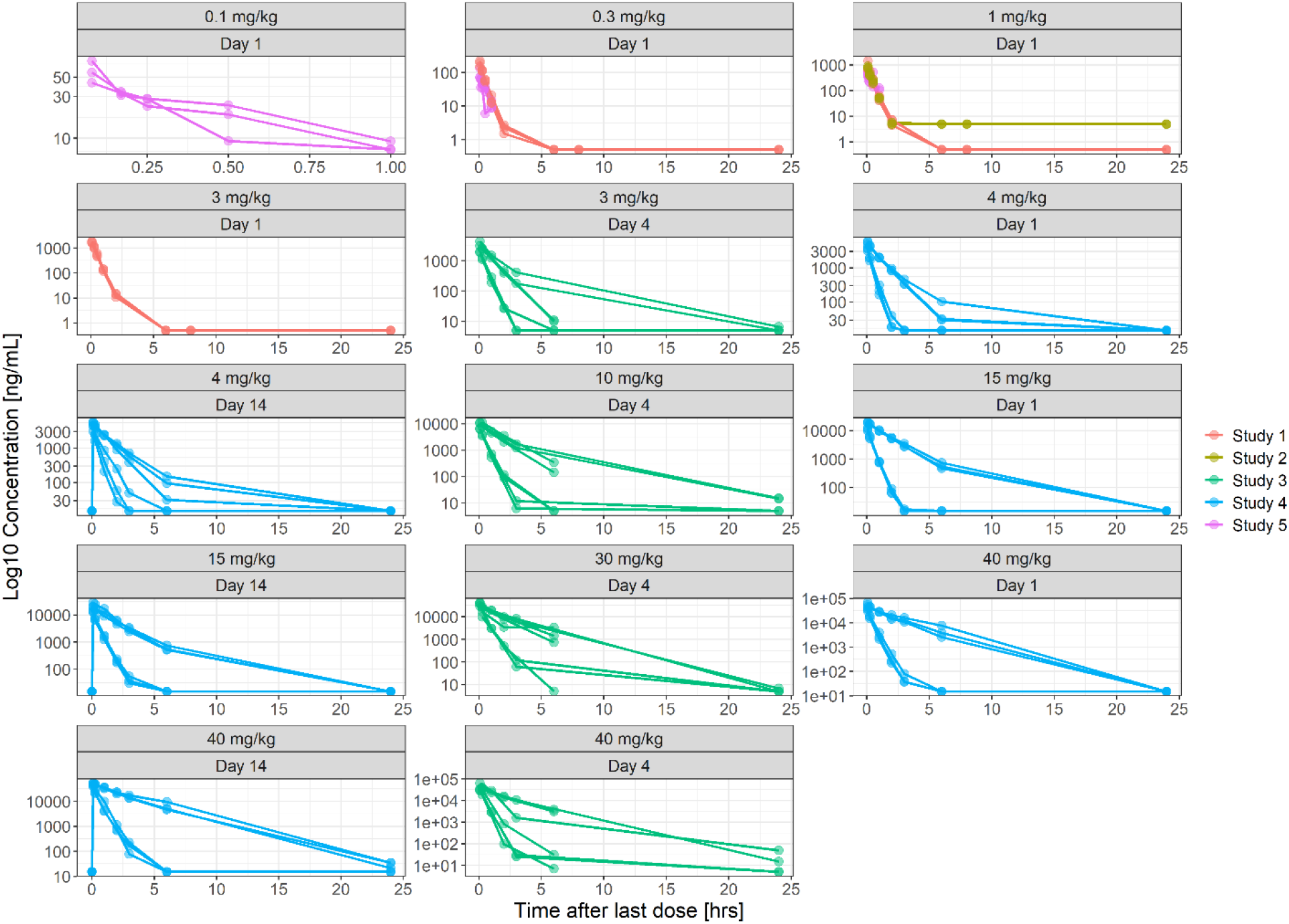
PK data in rats after administration of VC108 at time 0, stratified by administered dose and day of collection. Concentrations are shown on a logarithmic scale and colors are based on the study from which they were collected. Observations at time 0 represent pre-dose samples after repeated dose (e.g., at day 14).

In addition, 174 Sprague Dawley rats of both sexes were involved in the PD study, with measurements collected from distinct groups of animals which followed three alternative experimental scenarios differing by administration time of either VC108 or vehicle [22]. Control rats not receiving any drug or vehicle were also evaluated in groups 1 and 2. On average, PD (i.e. mpO_2_) measurements included 4 time-points per animal. PD data are summarized in Table 2 and Figure 2, respectively.

**Table 2.** PD data summary. For each experimental group considered in the PD study, the specific time of VC108 administration, sampling schedule, tested regimens, animal health status and number per sex are reported.

| Exp. Group | Administration timing | Sampling schedule (minutes)* | Regimen | Animal health status | Animal number per sex |
| --- | --- | --- | --- | --- | --- |
| 1 | 15 minutes before occlusion | 0, 30, 45<sup>§</sup>, 65, 105<br>0, 10, 30, 65, 105<br>0, 10, 45 or 30 and 65, 105 | Controls<br>Vehicles<br>0.5 mg/kg SD | Acute Myocardial Infarction | 13 M<br>6 M, 5 F<br>15 M, 12 F |
| 2 | 15 minutes after occlusion | 0, 20, 55, 95<br>0, 20, 35<sup>§</sup>, 55, 95 | Controls<br>0.5 mg/kg SD | Acute Myocardial Infarction | 12 M, 2 F<br>6 M, 7 F |
| 3 | 5 minutes before reperfusion | 0, 35, 60, 95<br>0, 35, 95<br>0, 35, 95<br>0, 35, 95<br>0, 35, 95<br>0, 20, 55, 95 | Vehicles<br>0.1 mg/kg SD<br>0.3 mg/kg SD<br>0.5 mg/kg SD<br>1 mg/kg SD<br>5 mg/kg SD | Acute Myocardial Infarction | 17 M, 13 F<br>7 M, 8 F<br>5 M, 6 F<br>12 M, 14 F<br>6 M, 6 F<br>1 M, 1 F |

\* Notes: Time 0 (i.e. baseline) is the time of the first collected mpO<sub>2</sub> value.
<sup>§</sup> PD sample collected only for one subject of the group.

**Fig. 2.**
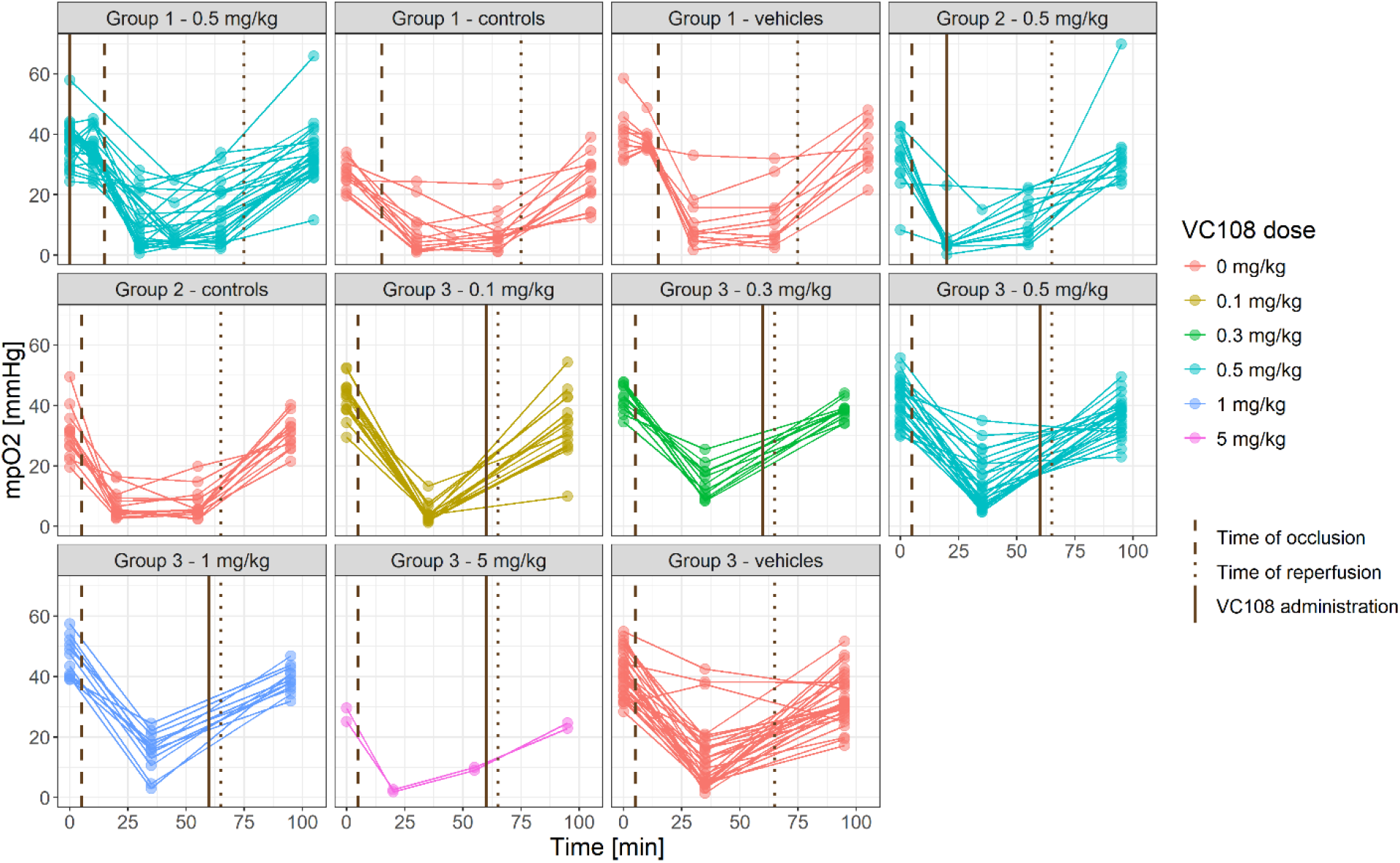
PD data in AMI rats model stratified by experimental group and administered dosage. Vertical lines indicate time of occlusion (dashed), time of reperfusion (dotted) and time of VC108 administration (solid).

Animal sex and body weight were recorded and investigated as possible covariates for the PK/PD model. VC108 was administered as single doses ranging from 0.1 to 40 mg/kg for PK assessment and 0.1 to 5 mg/kg for PD response characterization. In the PK study using the AMI rat model where drug concentrations were measured in whole blood rather than plasma, the experimentally determined blood-to-plasma (B/P) ratio of 0.905 was applied to convert these measurements to plasma concentrations, ensuring consistency across studies. The lower limit of quantification (LLOQ) for VC108 in plasma varies across studies between 0.5 ng/mL and 15 ng/mL. The LLOQ in blood was 2.5 ng/mL.

### PopPK and popPK/PD model development

During the development of the VC108 population pharmacokinetic (PopPK) model, both one- and two-compartment structures with linear and nonlinear elimination were evaluated. Then, a popPK/PD model was constructed by linking population parameter estimates from the popPK analysis with measured mpO_2_ levels. To the best of our knowledge, no models describing mpO_2_ during AMI have been previously reported. Therefore, we based the popPK/PD framework on the principles of indirect-response models that are well suited to variables at steady state that exhibit transient deviation following perturbation before returning to baseline [36,37]. This modelling strategy has been successfully applied in several studies describing drug effects on physiological pressure dynamics [38-41], providing theoretical support for our approach.

The development of the indirect effect popPK/PD model was designed as a two-step process. In the first step, mpO_2_ dynamics during AMI were characterized using only vehicle and control data, assuming vehicle administration had negligible effects on mpO_2_ flux; accordingly, subjects not treated with VC108 were hereafter referred to as “controls”. In the second step, a Bayesian approach was used to incorporate information from the first step and extend the baseline model to describe mpO_2_ dynamics under VC108 treatment.

PopPK modeling was performed assuming lognormal distributions for all parameters. Efficacy-related parameters were assumed to be lognormally distributed when physiologically constrained to positive values, or logitnormally distributed when constrained to values between 0 and 1.

### Model selection and evaluation

Model selection was guided by the corrected Bayesian Information Criterion (BICc), goodness-of-fit (GoF) diagnostics, and biological plausibility. Inter-Individual Variability (IIV) in model parameters was quantified, and potential covariates, i.e. body weight (BW), sex, and nominal dose, were evaluated through a manual stepwise covariate modeling approach. Once the covariate structure was established, the Conditional Sampling for Stepwise Approach based on Correlation tests (COSSAC) algorithm was employed to explore alternative covariate-model structures automatically [42]. Model performance was ultimately assessed using GoF plots and prediction-corrected visual predictive checks (pcVPC).

All popPK and popPK/PD analyses were conducted using a nonlinear mixed-effects modeling framework in Monolix 2024R1 (Lixoft).

### PopPK and popPK/PD simulations

To characterize the behavior of VC108 under different experimental scenarios and to evaluate the potential for administering an efficacious yet safe dose, a simulation-based approach was applied. An *in-silico* population of 500 rats (250 males and 250 females) was generated, with BW randomly sampled by a normal distribution with different means based on animal sex (males: mean 0.35 kg, SD 0.025 kg; females: mean 0.26 kg, SD 0.025 kg). Population typical values were assumed for PK parameters, while estimated inter-subject variability was considered for PD parameters; moreover, the impact of covariates selected in the popPK/PD model was considered for both PK and PD parameters. VC108 dosing was then simulated in this population as a bolus injection of 0 (controls), 0.1, 0.2, 0.3, 0.4, 0.5, 1, and 2.5 mg/kg. For each tested dose, mpO_2_ values over time were simulated considering three different administration schedules: (i) 15 minutes before coronary occlusion, (ii) 15 minutes after coronary occlusion, and (iii) 5 minutes before reperfusion, in alignment with the considered PD study (described in Table 2).

Simulated dose-response relationship was then analyzed by graphically comparing population median mpO_2_ variation relative to baseline (i.e., the median computed on individual mpO_2_ levels normalized to the value at time 0) across increasing doses. To quantify the effect of different VC108 dose levels on potential permanent damage, i.e. a reduction in mpO_2_ at steady state after AMI, the percentage of animals showing damage below certain thresholds was computed. Damage thresholds of 5%, 10%, 15% and 20% were investigated.

All popPK and popPK/PD simulations were conducted using the R package RsSimulx (R version 4.5.1).

## Results

### PopPK modeling

PK profiles were best described by a two-compartment model with linear elimination. Residual unexplained variability (RUV) was characterized using a proportional error model. The resulting model parameter estimates along with their precision are summarized in Table 3, where V1 is the volume of the central compartment, V2 the volume of the peripheral compartment, Cl the drug clearance, Q the inter-compartmental clearance and b the coefficient of the proportional residual error.

**Table 3.** PopPK model results. Estimated values for structural pharmacokinetic parameters are reported with corresponding relative standard errors (RSE) shown in square brackets and coefficient of variation (CV) of the inter-individual variability.

| Parameter |  | Covariate coefficients (Value [RSE %]) |  |  | Inter-Individual Variability |  |
| --- | --- | --- | --- | --- | --- | --- |
| Name [unit] | Value [RSE %] | Sex (= Male) | log BW | Nominal Dose | Value [RSE %] | CV % |
| Cl [L/h] | 0.23 [9.2] | 1.6 [7.5] | 0.99 [28.0] | -0.012 [19.1] | 0.24 [10.6] | 24.33 |
| V1 [L] | 0.41 [2.5] | - | 2.07 [5.8] | - | 0.11 [21.2] | 11.23 |
| Q [L/h] | 0.0035 [60.5] | - | - | - | 1.35 [23.1] | > 100 |
| V2 [L] | 0.091 [31.4] | - | - | - | 0.93 [23.0] | > 100 |
| b [-] | 0.45 [4.1] | - | - | - | - | - |

As a result of covariate assessment, clearance was found to be influenced by sex, BW, and nominal dose. At the doses studied, female rats exhibited greater exposure to VC108 compared to males, consistent with their lower clearance (Cl in females is 0.23 L/h versus 1.16 L/h in males). Body weight modeled using a logarithmic transformation normalized to the population median value 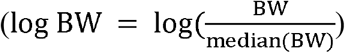 showed a negative association with clearance, meaning that heavier animals exhibited reduced clearance. Nominal dose also emerged as a significant covariate, with clearance decreasing at higher doses. Although alternative nonlinear models (e.g., Michaelis–Menten, Hill) were evaluated, they yielded unstable and unsatisfactory parameter estimates. With a β-coefficient of −0.012 on the log scale, meaningful changes in the clearance due to the administered amount per kg occur only at very high doses, which are outside of the range of interest for efficacy [22]. To further assess the impact of dose on clearance, an exploratory analysis is reported in the Supplementary Material (Figure S8). Finally, high coefficient of variation (CV>100%) resulted on the IIV of Q and V2; however, all the available covariates showed a statistically significant relationship with these two parameters.

PK model diagnostics, including individual fittings, GoF and pcVPC plots, are reported in the Supplementary Material (Figures S1 to S7).

### Indirect effect model for mpO_2_ in AMI model without treatment

The initial structural model was adapted from a typical indirect effect model structure, considering zero-order mpO_2_ input and first-order consumption. To better capture the dynamics observed after release of coronary occlusion, an overshoot (OS) component of mpO_2_ was incorporated, as it is physiologically plausible [6]. The resulting model, used to describe data derived from controls, is presented in Eq. (1).

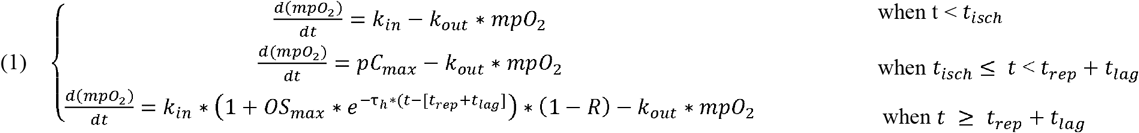

Specifically, *t*_*isch*_ denotes the time at which ischemia is induced (left coronary artery ligation), and *t*_*rep*_ the time at which the occlusion is released. *k*_*in*_ represents the mpO_2_ influx delivered through the left main coronary artery, while *k*_*out*_ describes myocardial O_2_ consumption. While before ischemia the left coronary artery provided full O_2_ flux to the area at risk, no O_2_ is provided to this region from the left coronary artery during ischemia (*k*_*in*_ = 0). *pC*_*max*_ accounts for cBF that sustains a residual O_2_ supply during occlusion.

After reperfusion, O_2_ starts flowing back to the myocardium from the left coronary artery; however, damage already caused by preceding ischemia is represented by the *R* parameter. *OS*_*max*_ reflects the maximal increase in O_2_ influx after reperfusion, and *τ*_*h*_ is the time constant governing its exponential decline toward baseline. Finally, *t*_*lag*_ denotes the delay that can occur between reperfusion (*t*_*rep*_) and the onset of the observed rise in O_2_ influx.

This framework captures mpO_2_ dynamics across three phases of the AMI: (i) at baseline, before coronary occlusion, (ii) the ischemic period during coronary occlusion, and (iii) the reperfusion phase after release of coronary occlusion. Since mpO_2_ measurements were sparse, a priori information was introduced for parameters governing the overshoot component, guided by literature [10,15,18]. Specifically, *OS*_*max*_, the maximal overshoot relative to baseline (expressed as a fraction), was assigned a lognormal distribution with mean 0.75 and standard deviation 1 as prior for Maximum A Posteriori (MAP) estimation. Similarly, the time constant τ_*h*_, describing the exponential decline of the overshoot toward steady state, was assigned a lognormal prior with mean 0.15 and standard deviation 1 as prior for MAP algorithm. This choice assumes that the overshoot dissipates within roughly 0.7 hours (τ_h_ ≈ 0.7 h / 4.5 h). No measurements were made after this period.

At steady state, baseline mpO_2_ is defined by the ratio pO_2,0_ *k*_*in*_/*k*_*out*_. To facilitate parameter estimation, mpO_2_ was made explicit in the model that was subsequently reparametrized. Finally, RUV was modeled using both constant (*a*) and proportional (*b*) error components. Model results are summarized in Table 4.

**Table 4.** PopPD model results in non-treated AMI rats model. Estimated values for structural parameters are shown with their corresponding RSE in square brackets and CV of the inter-individual variability.

| Parameter |  | Covariate coefficients (Value [RSE %]) |  |  | Inter-Individual Variability |  |
| --- | --- | --- | --- | --- | --- | --- |
| Name [unit] | Value [RSE %] | Sex (= Male) | log BW | Nominal Dose | Value [RSE %] | CV % |
| $\text{pO}_{2,0}$ [mmHg] | 35.39 [2.7] | - | - | - | 0.17 [15.9] | 17.16 |
| $k_{out}$ [1/min] | 0.52 [14.7] | - | - | - | 0.36 [24.9] | 36.71 |
| $pC_{max}$ [mmHg/min] | 4.14 [16.6] | - | - | - | 0.63 [16.2] | 69.55 |
| $R$ [-] | 0.16 [11.7] | - | - | - | 0.56 [24.8] | 45.11 |
| $t_{lag}$ [min] | 0 [Fixed] | - | - | - | - | - |
| $OS_{max}$ [-] | 0.57 [46.4] | - | - | - | 1.1 [35.5] | > 100 |
| $\tau_h$ [-] | 0.15 [43.0] | - | - | - | 0.8 [42.2] | 95.27 |
| $a$ [mmHg] | 1.4 [24.0] | - | - | - | - | - |
| $b$ [-] | 0.14 [16.5] | - | - | - | - | - |

All parameters were estimated with good precision, as demonstrated by the relative standard error (RSE) that are all lower than 50%. No delay between reperfusion and the onset of the rise in O_2_ influx was detected: indeed, if a *t*_*lag*_ was included into the model, its estimate resulted close to 0. For this reason, this parameter was finally fixed at 0, with a consequent improvement in model performance (i.e. lower BICc and better precision of the estimates).

### PopPK/PD indirect effect model for mpO_2_ in AMI model with VC108 treatment

After estimating the controls-derived model, the second step was to fit the model to the overall population including VC108-treated animals. The effect of VC108 was tested on parameters governing O_2_ influx, consistent with its role as a GPR39 blocker expected to enhance microvascular perfusion and thereby improve myocardial O_2_ delivery during AMI [21,22]. Prior information derived from the estimates obtained in the previous step were adopted for non-drug related parameters and used to perform a Bayesian estimation process. The model that best fits the data is described in Eq. (2).

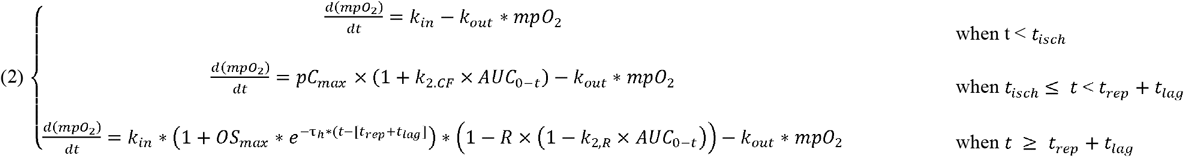

The model incorporates augmented cBF during coronary occlusion and a reduced extent of permanent damage, both expressed as functions of cumulative drug exposure, with magnitudes proportional to parameters *k*_*2.CF*_ and *k*_*2,R*_, respectively.

As shown in Table 5, an additional parameter, *λ*, in the parameters list. This parameter defines the shape of the Box–Cox transformation applied to mpO_2,0_ to better capture the baseline distribution of mpO_2_ in individual animals. When considering all data combined (with controls-only subjects accounting for 61 of the total of 174), the distribution of mpO_2,0_ was more consistent with a Box–Cox distribution [43] than with a lognormal one. This behavior was not evident in the controls data and, therefore, the parameter was incorporated only at this stage. Notably, assuming a lognormal distribution for this parameter resulted in a higher BICc value. In addition, a significant positive correlation (r=0.52, p-value<0.001) was found between *pC*_*max*_ and _mpO_η_2,0_ parameters and thus considered in the model. All parameters were estimated with good precision, achieving RSE of approximately ≤ 60%. The model adequately characterized the mpO_2_ dynamics, as demonstrated by the pcVPC at the population level and by the individual fits presented in Figures 3 and 4, respectively. In Figure 4, only a representative subset of individuals covering different administration times and regimens is shown, whereas the full set of individual fits for the analyzed population, as well as the GoF plot of PD data, is provided in the Supplementary Material (Figures S9 to S12).

**Table 5.** PopPK/PD model results in AMI rats model. Estimated values for structural parameters are shown with their corresponding RSE in square brackets and CV of the inter-individual variability.

| Parameter |  | Covariate coefficients (Value [RSE %]) |  |  | Inter-Individual Variability |  |
| --- | --- | --- | --- | --- | --- | --- |
| Name [unit] | Value [RSE %] | Sex (= Male) | log BW | Nominal Dose | Value [RSE %] | CV % |
| $pO_{2,0}$ [mmHg] | 39.06 [1.9] | - | - | - | - | - |
| $\eta_{po_{2,0}}$ [-] | 0 [Fixed] | - | - | - | 0.13 [22.3] | > 100 |
| $\lambda$ [-] | -4.3 [34.1] | - | - | - | - | - |
| $k_{out}$ [1/min] | 0.69 [11.0] | - | 0.66 [44.0] | - | 0.53 [13.9] | 56.43 |
| $pC_{max}$ [mmHg/min] | 5.32 [11.6] | - | - | - | 0.44 [25.5] | 46.55 |
| $R$ [-] | 0.12 [11.6] | - | - | - | 0.81 [12.1] | 67.45 |
| $t_{lag}$ [min] | 0 [Fixed] | - | - | - | - | - |
| $OS_{max}$ [-] | 0.36 [56.0] | - | - | - | 1.49 [21.1] | > 100 |
| $\tau_h$ [min] | 0.23 [30.1] | - | - | - | 0.72 [15.4] | 82.49 |
| $k_{2,CF}$ [mL/(mg*min)] | 0.06 [38.2] | - | - | - | 1.06 [27.0] | > 100 |
| $k_{2,R}$<br>[mL/(mg*min)] | 0.004<br>[61.3] | - | - | - | 1.27<br>[34.0] | > 100 |
| corr_<br>$pC_{max-\eta_{pO_{2,0}}}$ | 0.52<br>[31.3] | - | - | - | - | - |
| a [mmHg] | 1.88<br>[15.5] | - | - | - | - | - |
| b [-] | 0.11<br>[21.7] | - | - | - | - | - |

**Fig. 3.**
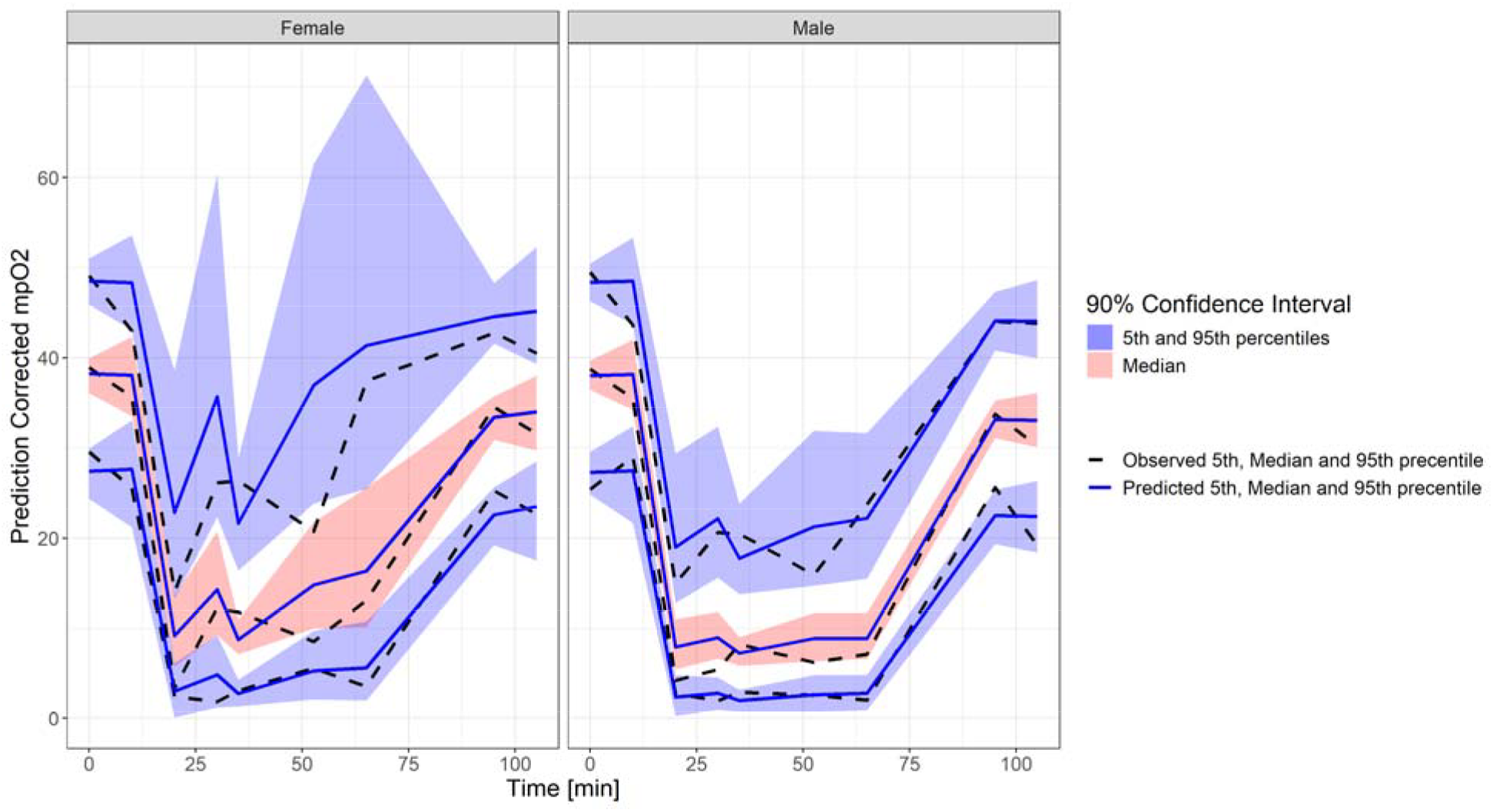
pcVPC of the developed PK/PD model, stratified by sex. Black dashed lines represent observed 5th, 50th (median) and 95th percentiles, while blue solid lines represent predicted 5th, 50th (median) and 95th percentiles. Filled areas indicate 90% confidence interval, in particular the red one represents the median while the blue ones represent 5th and 95th percentiles, respectively.

**Fig. 4.**
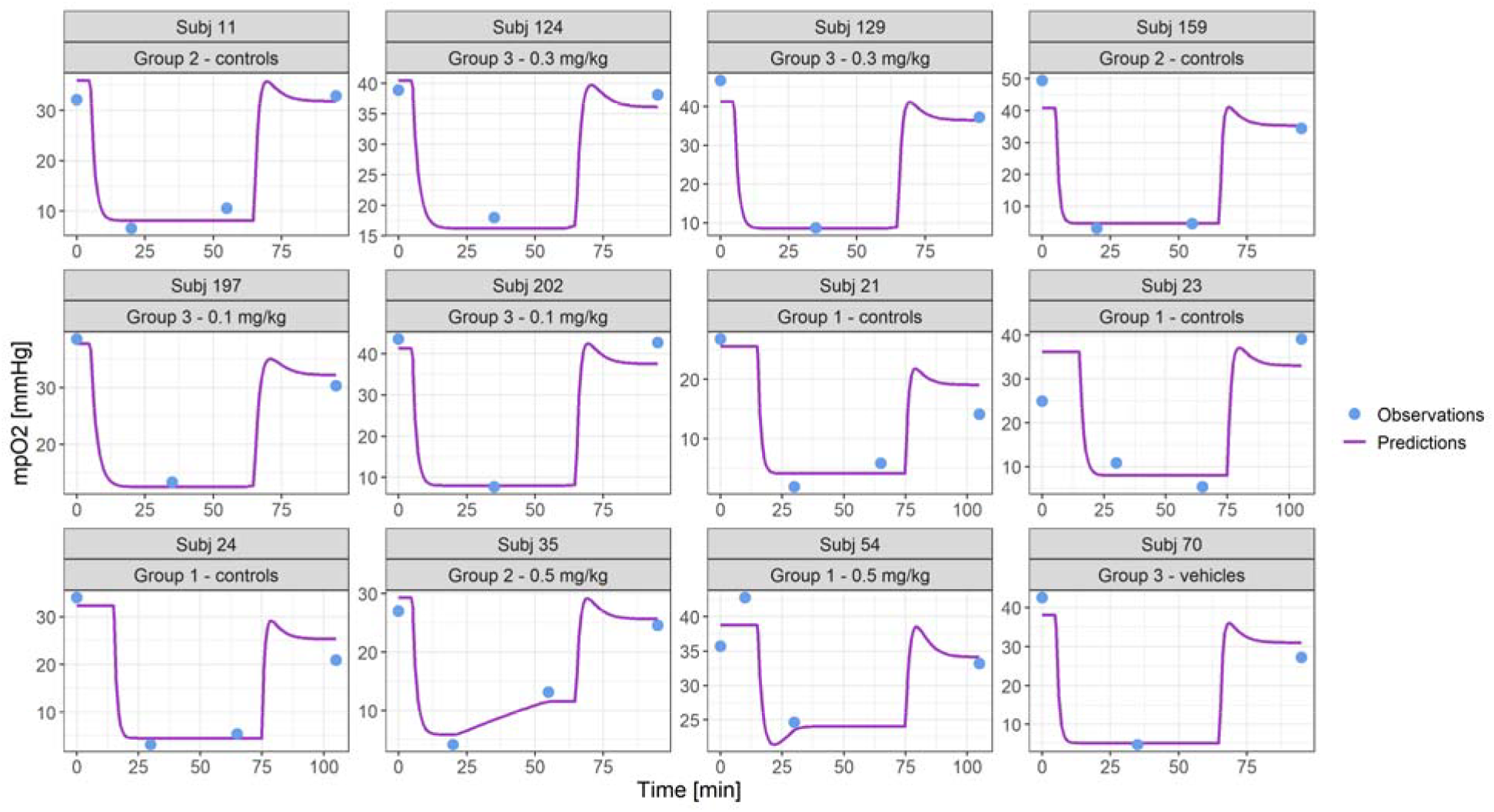
Model individual fittings of PD data in a representative subset of individuals covering different administration times and regimens. Blue points are the observations while purple lines are the model predictions.

### Simulations of mpO_2_ profile with different dosing regimens

A total of 500 virtual subjects (250 males and 250 females) were simulated for each dose and administration scenario. The model predicted a clear dose–response relationship, as shown in Figure 5, where the median mpO_2_ variation relative to the baseline is compared across increasing doses and controls data.

**Fig. 5.**
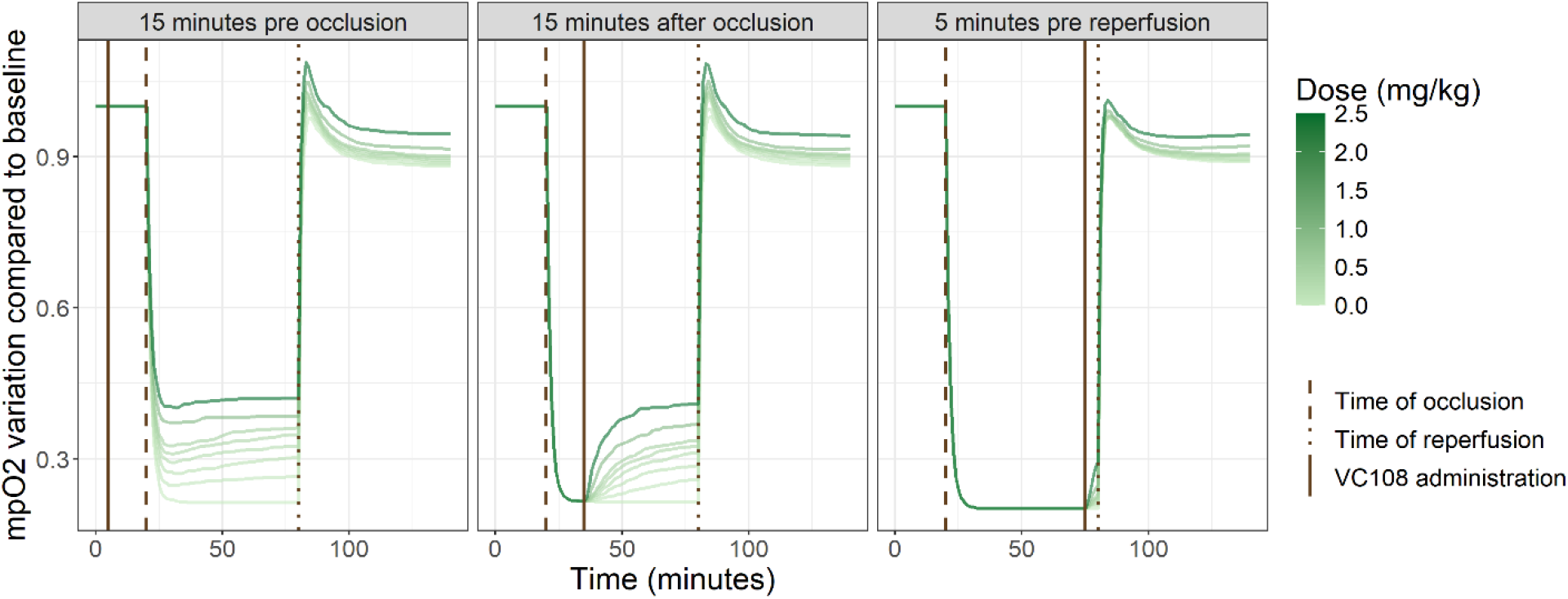
Median mpO_2_ variation relative to baseline of the simulated population treated with increasing doses and considering three different administration scenarios. Vertical lines indicate time of occlusion (dashed), time of reperfusion (dotted) and time of VC108 administration (solid).

In the two scenarios where drug was administered 15 minutes before and 15 minutes after coronary occlusion, VC108 enhanced cBF, providing increased O_2_ delivery to the area at risk during coronary occlusion. In the scenario where VC108 was administered 5 minutes before reperfusion, an increase in cBF is still observed, but it is limited due to the shorter drug exposure time. When comparing the reperfusion induced overshoot in mpO_2_ levels for a given dose, the peak is lower when drug is given 5 minutes before reperfusion. Because drug effect is dependent on the overall exposure (AUC_0–t_), when VC108 is administered only 5 minutes before reperfusion, the resulting AUC_0-tlast_ (in median 12677 ng/mL□min for 0.5 mg/kg) is lower than the AUC_0-tlast_ reached when VC108 is administered either 15 minutes before or after coronary occlusion (in median 17646 ng/mL□min and 15915 ng/mL□min for 0.5 mg/kg). Consequently, the reduced effect observed in the latter scenario is associated with a shorter duration between VC108 administration and time of reperfusion, and thus a smaller increase of cBF during occlusion.

Since PK parameters markedly differ between males and females, results were stratified to highlight sex-specific differences (Figure 6). It is evident that females exhibit a greater VC108 effect. Interestingly, no sex-related covariates were identified for any PD parameters. Therefore, according to the model, this difference can be attributed to variations in drug exposure rather than to differences in the mechanism of action of VC108 between males and females.

**Fig. 6.**
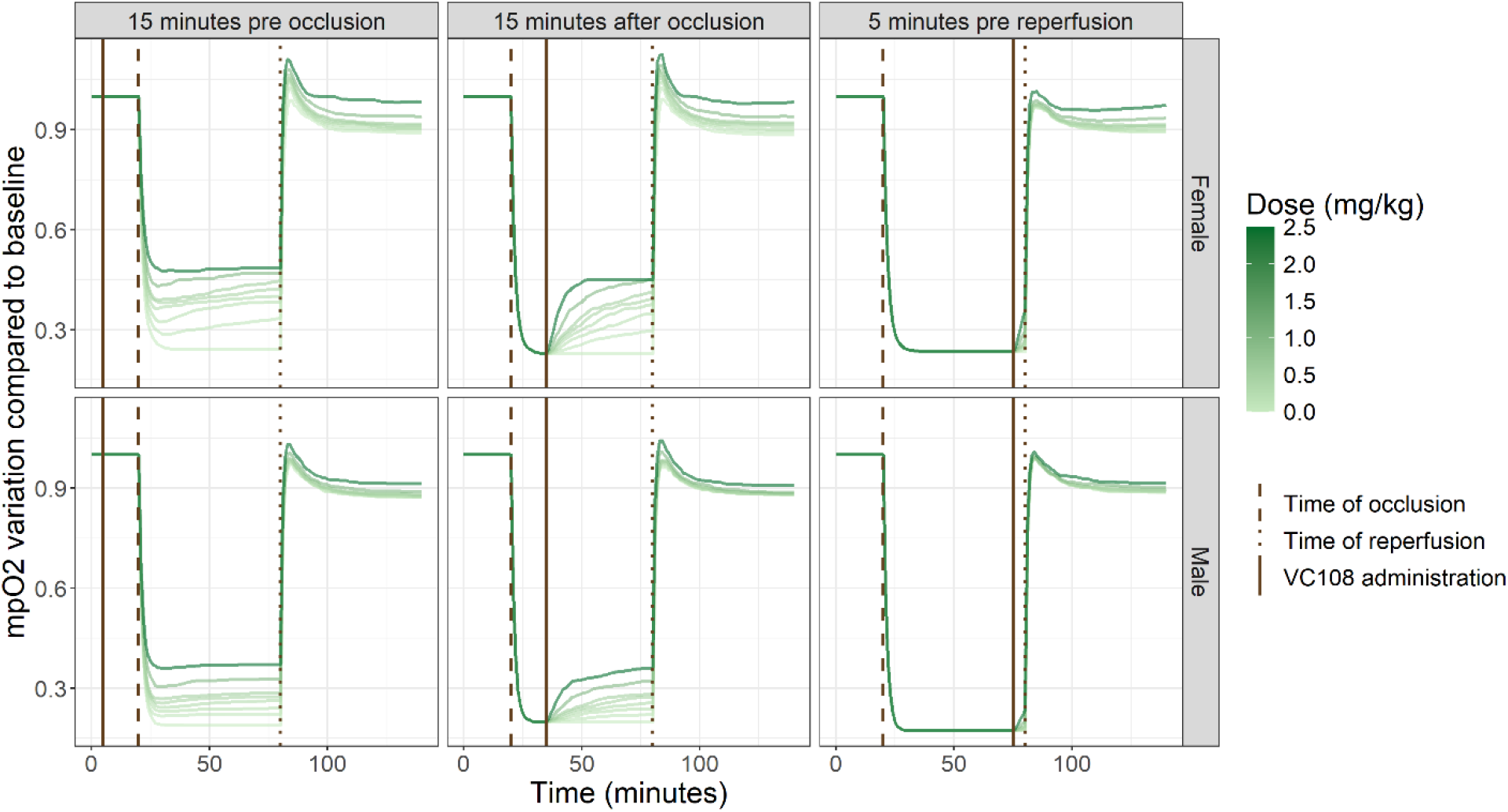
Median mpO_2_ variation relative to baseline of the simulated population treated with increasing doses and considering three different administration scenarios, stratifying by sex. Vertical lines indicate time of occlusion (dashed), time of reperfusion (dotted) and time of VC108 administration (solid).

To further examine the simulations, focusing on the more clinically likely scenario (i.e., VC108 administered 5 minutes before reperfusion), the percentage of simulated individuals with permanent damage, i.e. an irreversible reduction in mpO_2_ at steady state after AMI, is computed. The results are shown in Figure 7 for different administered doses and damage thresholds (5%, 10%, 15% and 20%) stratified by sex.

**Fig. 7.**
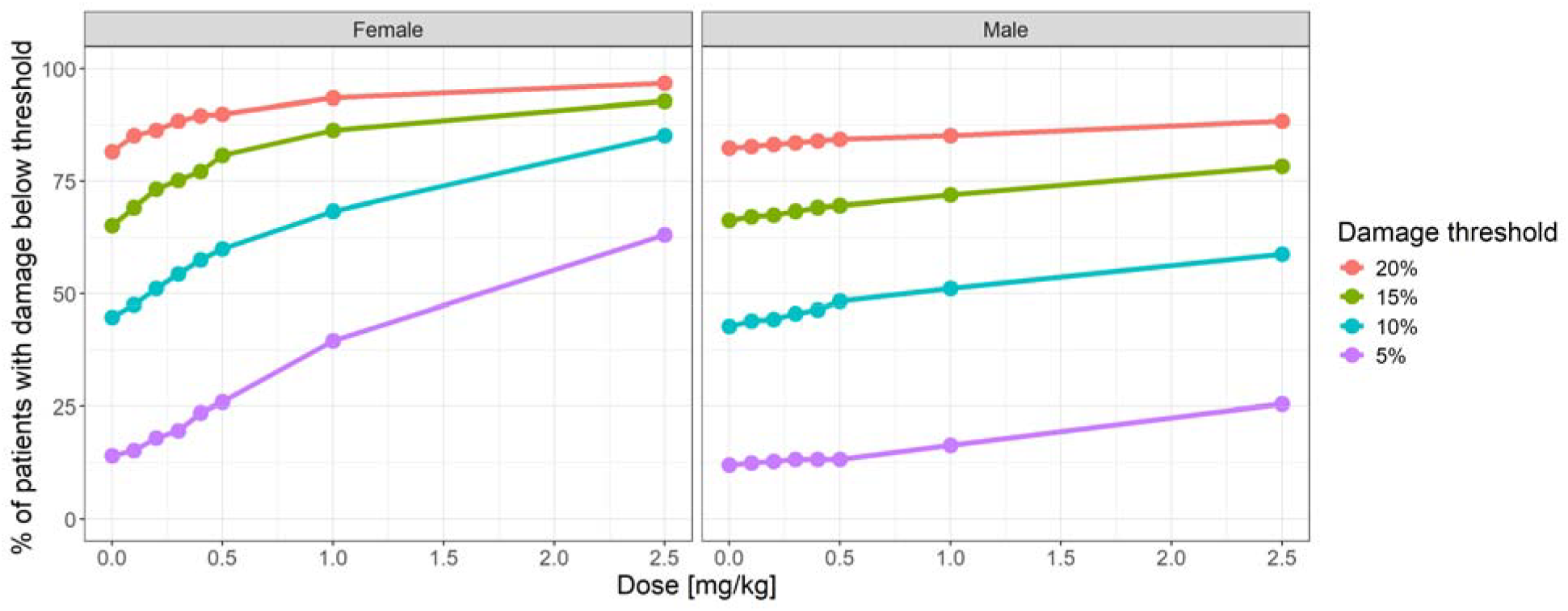
Percentage of patients with permanent damage below 5%, 10%, 15% and 20% for different doses administered 5 minutes before reperfusion. Curves are stratified by sex.

The probability of remaining below the predefined damage thresholds increases with dose in both sexes, with females consistently achieving greater protection. For females, the effect plateaus around 1□mg/kg when considering a higher permanent damage (i.e., 15% or 20%), whereas in males the effect increases more modestly across the entire dose range.

## Discussion

This study presents the first popPK/PD model describing the dynamics of mpO_2_ in AMI and its modulation by the GPR39 blocker, VC108. By linking drug exposure to time-dependent changes in tissue O_2_ availability, the model captures basal homeostasis, ischemic decline, and reperfusion overshoot, while quantifying the contribution of VC108 to cBF and functional recovery. This approach demonstrates the value of an indirect response structural model for characterizing physiological outcomes that are modified by the drug through its vascular mechanisms.

VC108 pharmacokinetic was well described by a two-compartment model with linear elimination and with BW, sex and nominal dose identified as significant covariates influencing clearance and central compartment volume (the latter only by BW). The finding that clearance decreased at higher nominal doses was unlikely to reflect true nonlinear elimination, but rather a drug-related reduction in clearance at very high exposures. Sex-related differences in clearance were particularly relevant with females exhibiting lower clearance and therefore higher systemic exposure compared to males at higher doses of VC108.

The structural popPK/PD model was then developed in two steps, beginning with control data (also comprising vehicle administration) to characterize mpO_2_ ischemia–reperfusion dynamics. The reperfusion overshoot was described using informative initial conditions for both magnitude and decay rate based on literature. Interestingly, even if the phenomena cannot be clearly seen from the data due to the sampling scheme of the study that prevented data collection in that specific timeframe, posterior estimates deviated remarkably (around 50%) from the prior means, suggesting that the available data, despite being relatively sparse, carried sufficient information to refine these parameters rather than simply reproduce initial assumptions. Another key feature of the model was the quantification of cBF during coronary occlusion, which accounted for approximately 20% of initial O_2_ flow (*pC*_*max*_ versus *k*_*out*_ * mp*O*_2,0_ terms of Eq. (1)). This estimate is physiologically plausible and consistent with the presence of residual perfusion within the area at risk in the rat model [6,44]. The finding that VC108 enhances cBF during ischemia supports critical contribution of the collateral circulation for sustaining myocardial oxygenation under ischemic conditions and constitutes a meaningful target for pharmacological modulation.

The exposure–response simulations also highlighted sex-related differences. Both sexes benefited from VC108, but females exhibited a greater response at the same given dose/kg compared to males. In males, the protective effect increased more gradually across the entire dose range, suggesting that higher exposures may be required to achieve comparable levels of protection. Importantly, the favorable tolerability profile of VC108 even at high doses in both sexes indicates that escalation in males may be feasible without major safety concerns. It should be noted, however, that the toxicology profile of VC108 is not addressed in this work, as it lies beyond the scope of the present PK/PD analysis. The observed sex difference was not explained by pharmacodynamic parameters but was fully attributable to sex-related differences in clearance and exposure.

To conclude, the study strengthens the rationale of using mpO_2_ as a pharmacodynamic endpoint because it represents a mechanistically relevant and relatively easy-to-measure biomarker that integrates the net balance between myocardial O_2_ supply and consumption. The modeling framework allowed us to (i) describe ischemia–reperfusion dynamics with biologically grounded parameters, (ii) quantify drug effects on cBF, and (iii) predict sex-specific outcomes in untested dose regimens. Beyond VC108, this framework illustrates how indirect response models can be applied to cardiovascular endpoints in the context of AMI, providing a broadly applicable approach for mechanistic PK/PD analysis in cardiovascular pharmacology.

Some limitations should also be acknowledged. First, the experimental design was not optimized for population PK/PD analysis. PK/PD data were obtained from separate sets of animals, precluding subject-level correlations. Second, the number of mpO_2_ measurements per animal was limited, requiring the use of informative priors and potentially introducing uncertainty in parameter estimation. Third, mpO_2_ tension was measured by injecting the oxyphor into the anterolateral myocardium, which does not provide information on the spatial distribution of blood flow. Consequently, we cannot determine the effects of cBF in the margins of the area at risk but only in its center. The fact that even the central ischemic zone was favorably affected by cBF suggests that the marginal regions are likely to be better affected because of the gradient of flow between the margins and the center. Despite these limitations, the developed model represents a powerful tool which allowed, for the first time, to well describe VC108 efficacy data and quantify its effect in untested dose levels.

## Conclusions

This study presents the first population PK/PD model of mpO_2_ in AMI rat model, integrating VC108 pharmacokinetics with ischemia–reperfusion dynamics. VC108 improved myocardial oxygenation primarily by enhancing cBF, leading to dose-dependent reductions in permanent damage. The model revealed sex- and BW–dependent clearance, with females achieving higher exposure and greater protection at equivalent doses. Importantly, sex differences were explained by pharmacokinetics rather than pharmacodynamics. Beyond its relevance to VC108 as a promising drug, this framework illustrates how PK/PD modeling can quantitatively disentangle blood flow, mpO_2_ balance, and drug action, providing a translational tool for cardiovascular pharmacology to inform the selection of an initial effective dose and further refine dosing strategies.

## Supporting information

https://www.dropbox.com/scl/fi/lybetbq431w6pgjxbsl99/Supplementary-Material_Ronchi-et-al.docx?rlkey=n84c82w7o2o2xd7han7cpkgz9&dl=0

## Supplemental Materials

Supplemental Text Supplemental Figures S1-S12

