## Supplementary material for "Impact of GPR39 Inhibition on Tissue Oxygen Tension in Acute Myocardial Infarction: Modeling Using an Indirect Response Pharmacokinetics/Pharmacodynamics Approach": https://www.dropbox.com/scl/fi/lybetbq431w6pgjxbsl99/Supplementary-Material_Ronchi-et-al.docx?rlkey=n84c82w7o2o2xd7han7cpkgz9&dl=0

**PK model diagnostics**

**Fig. S1** Model individual fittings of PK data in individuals belonging to Study 1. Blue points are the observations while purple lines are the model predictions.

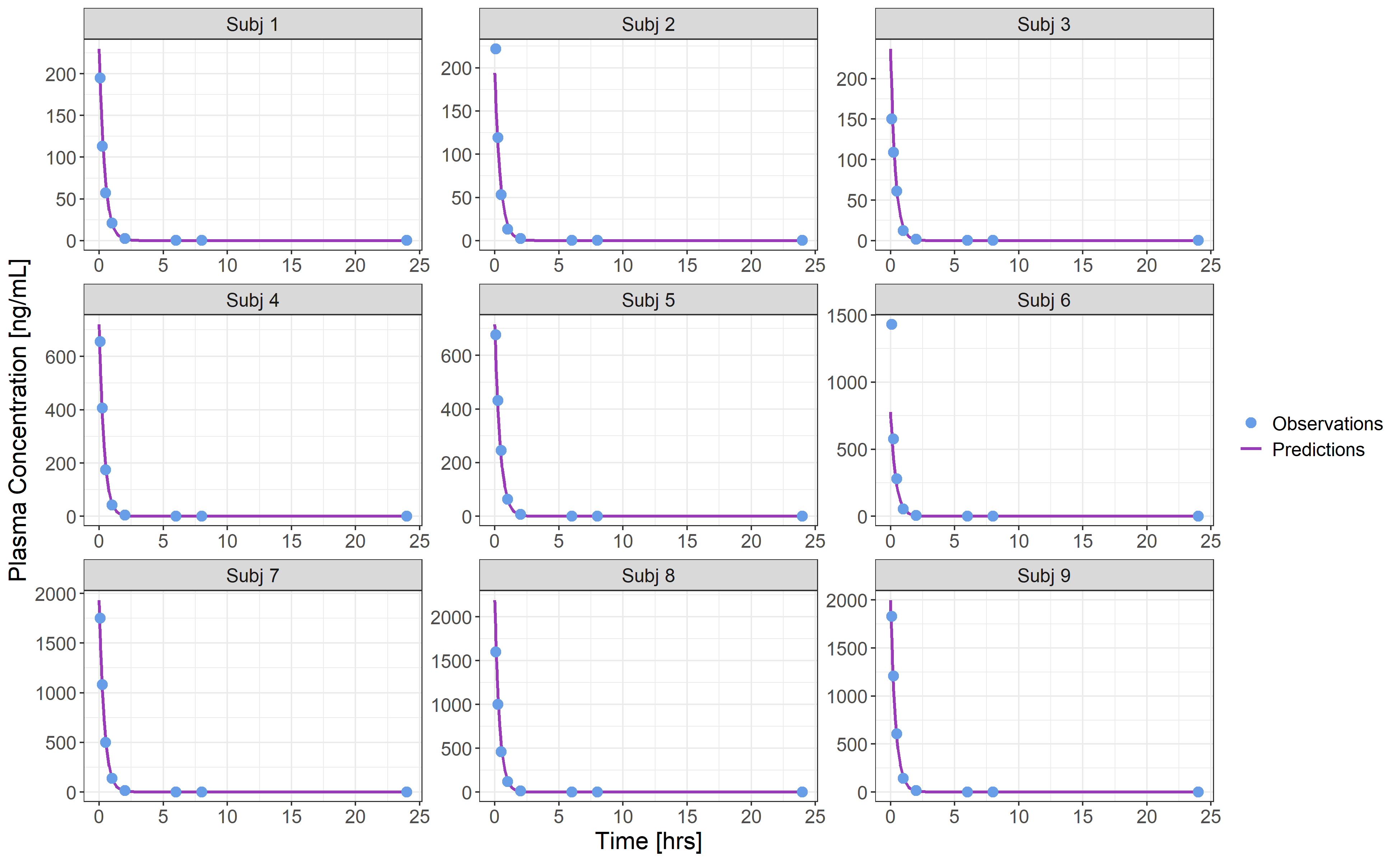

**Fig. S2** Model individual fittings of PK data in individuals belonging to Study 2. Blue points are the observations while purple lines are the model predictions.

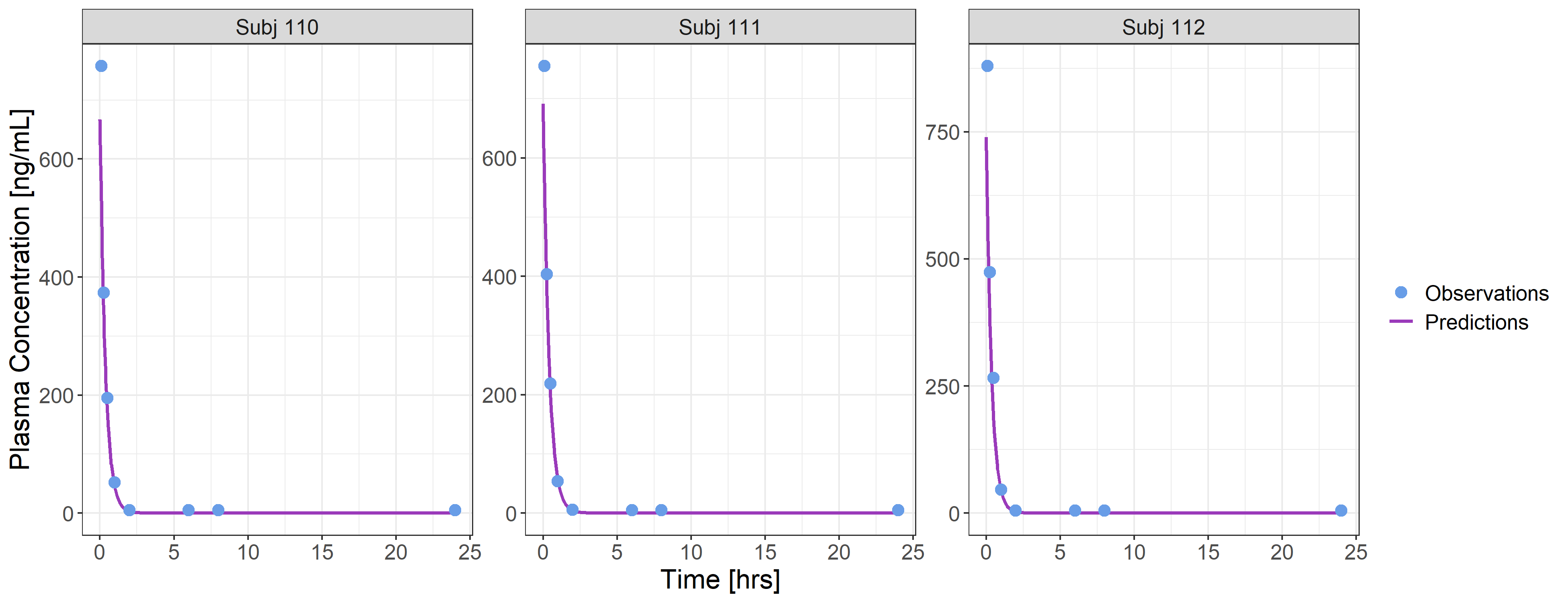

**Fig. S3** Model individual fittings of PK data in individuals belonging to Study 3. Blue points are the observations while purple lines are the model predictions.

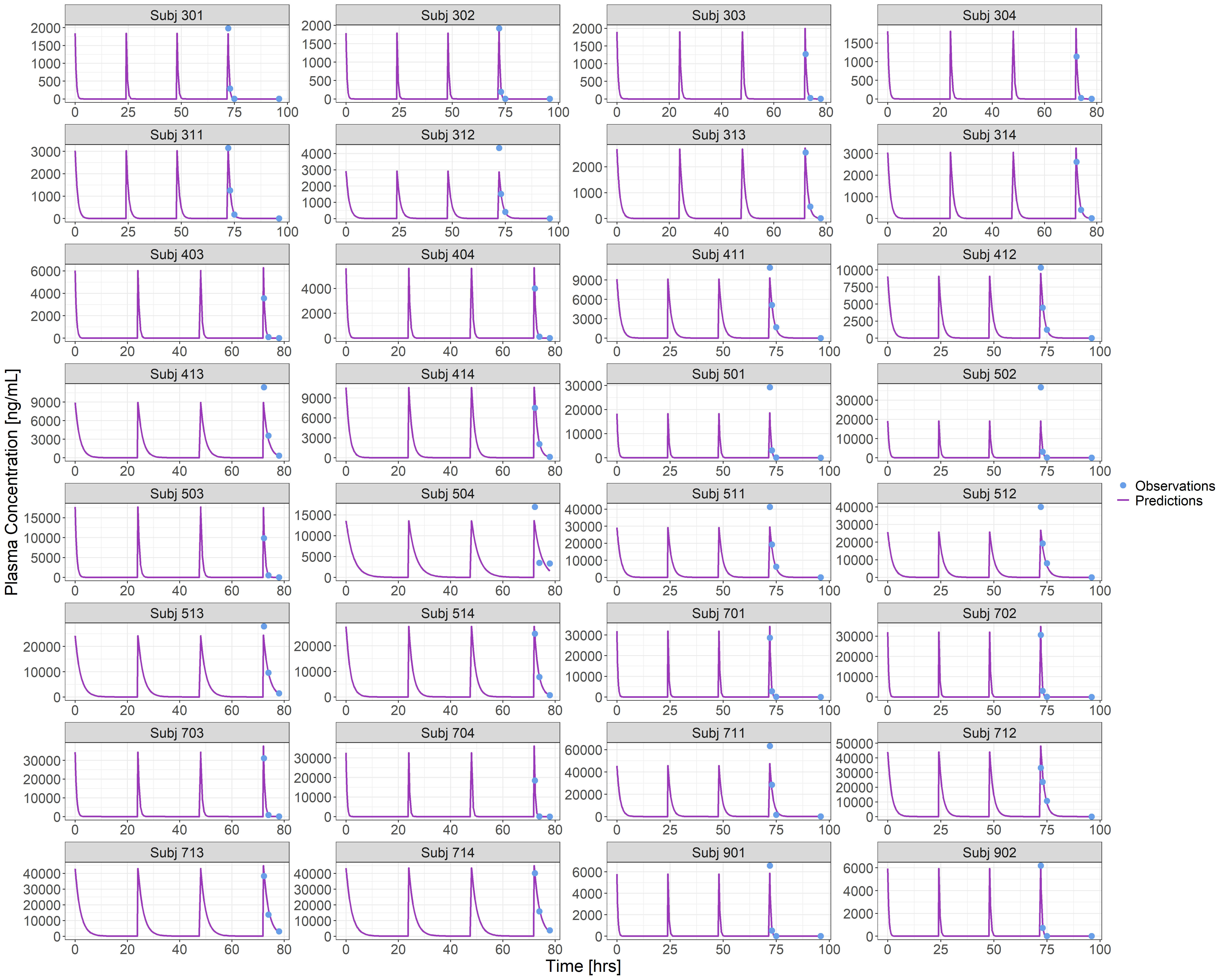

**Fig. S4** Model individual fittings of PK data in individuals belonging to Study 4. Blue points are the observations while purple lines are the model predictions.

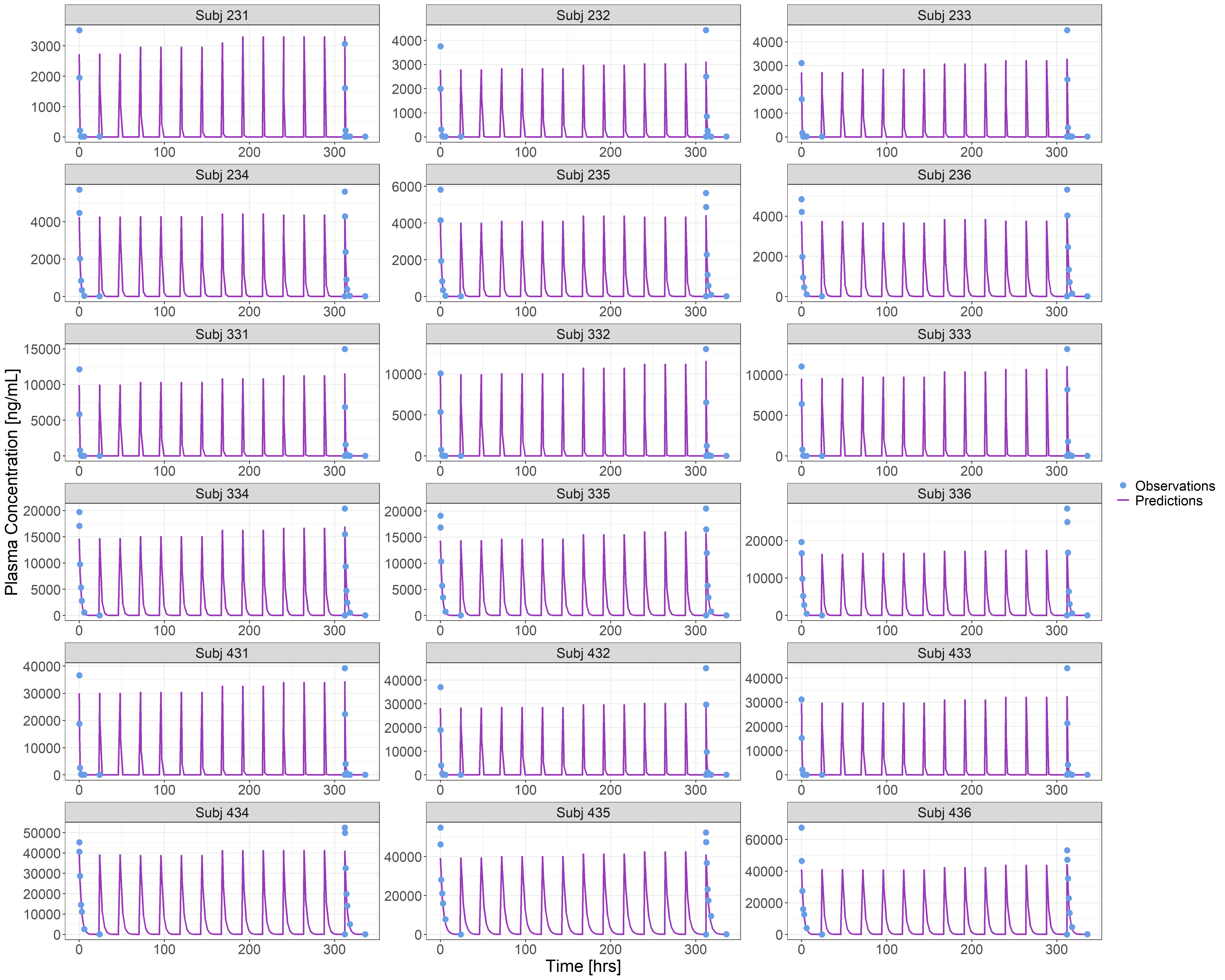

**Fig. S5** Model individual fittings of PK data in individuals belonging to Study 5. Blue points are the observations while purple lines are the model predictions.

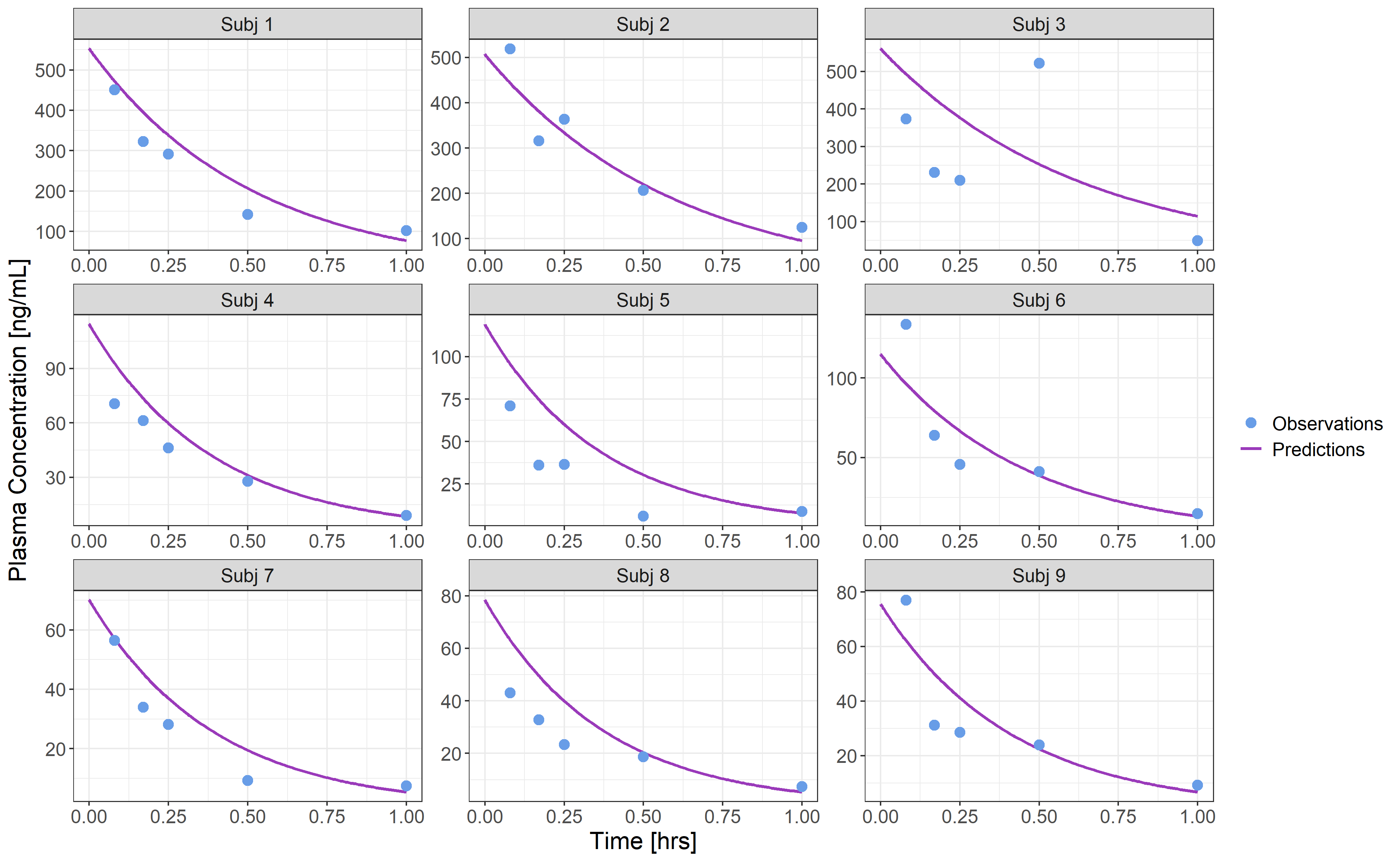

**Fig. S6** Model Goodness Of Fit plot of PK data (in Log scale). Population predictions are shown in the left box, while individual predictions are reported in the right box.

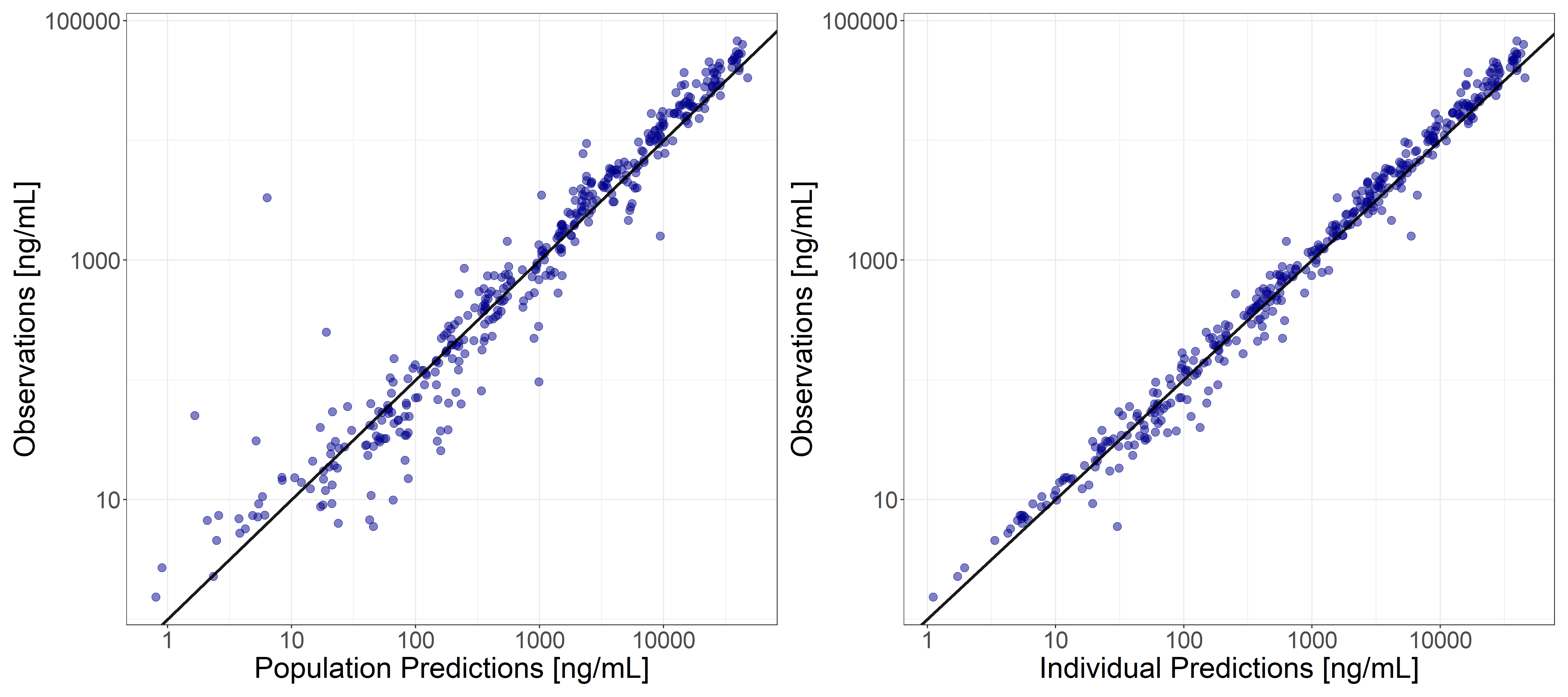

**Fig. S7** pcVPC of the developed PK model, stratified by sex. Black dashed lines represent observed 5^th^, 50^th^ (median) and 95^th^ percentiles, while blue solid lines represent predicted 5^th^, 50^th^ (median) and 95^th^ percentiles. Filled areas indicate 90% confidence interval, in particular the red one represents the median while the blue ones represent 5^th^ and 95^th^ percentiles, respectively.

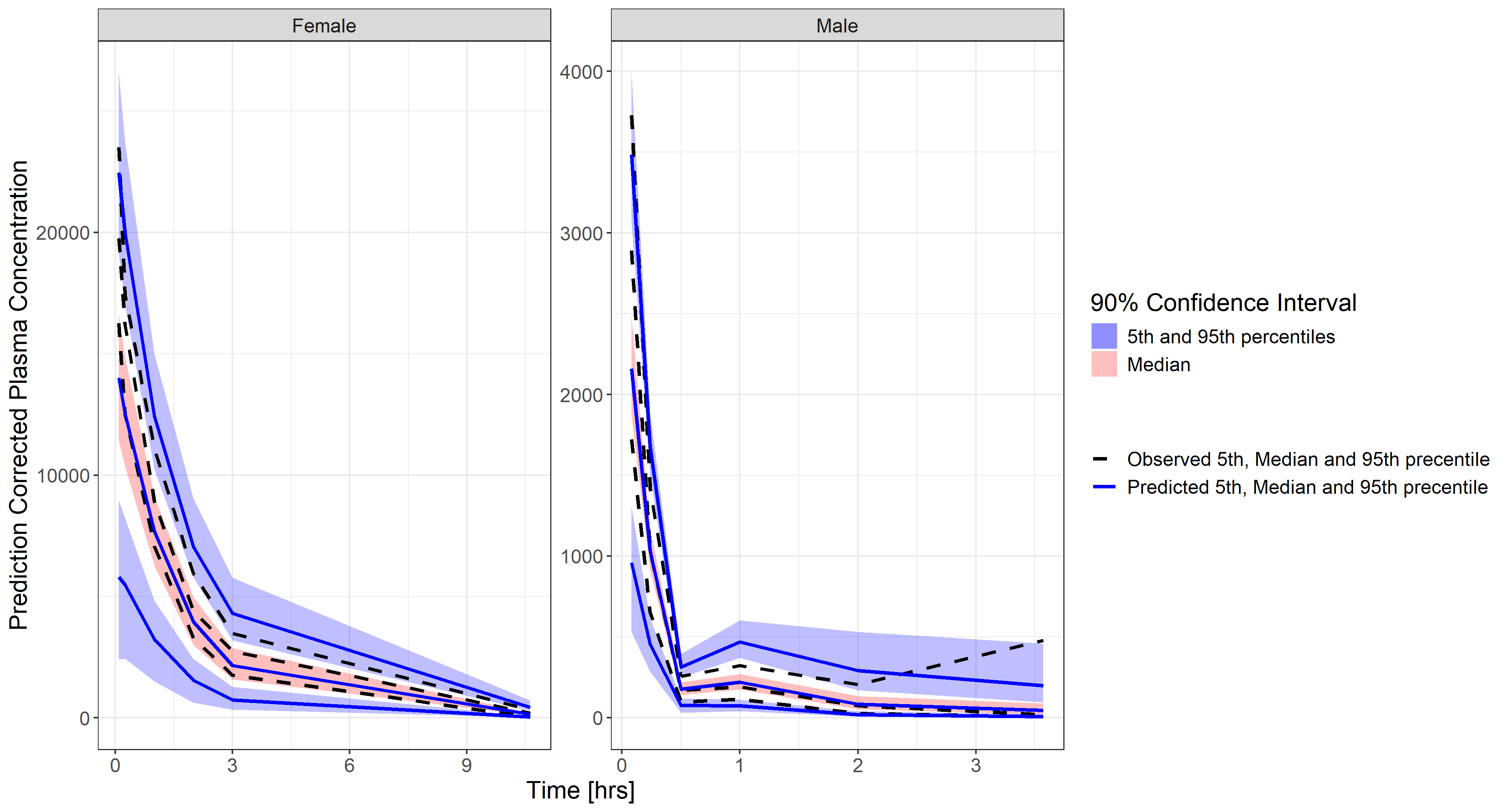

**Exploratory analysis to assess the impact of nominal dose on clearance**

Based on the developed PK model, nominal dose resulted in a significant covariate on clearance, with a beta coefficient of -0.012 on the log scale, meaning that meaningful changes in the clearance due to the administered amount per kg occur only at very high doses. To better assess this effect, the nominal dose was dichotomized into <30 mg/kg and ≥30 mg/kg, where 30 mg/kg was considered as cutting threshold being the one of the higher doses tested in the PK studies, and the resulting binary covariate was used in the PK model in place of the continuous one. This model provided a BICc only few points higher than the one obtained considering the dose as continuous covariate (6598.8 vs 6590.2, respectively), indicating that elimination remains practically linear at doses < 30 mg/kg, whereas clearance decreases at higher doses. A graphical comparison between the two models for a typical female subject, i.e. with BW equal to the population median BW, is reported in Figure S8. Overall, the results indicate that the apparent dose effect is unlikely to represent nonlinear pharmacokinetics but is more plausibly explained by a possible drug-related effect on clearance at elevated exposures.

**Fig. S8** Effect of nominal dose on clearance in a typical female subject under the continuous (black solid line) and dichotomous (red dashed line) covariate models. Points indicate tested doses and the vertical red line marks the dichotomization threshold (i.e. 30 mg/kg).

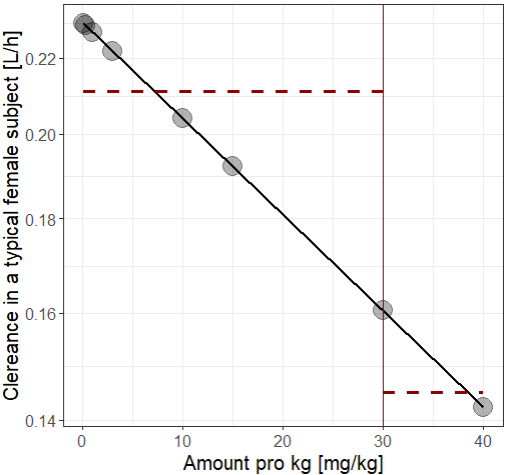

**PD model diagnostics**

**Fig. S9** Model individual fittings of PD data in individuals belonging to experimental group 1 (drug administration 15 minutes before occlusion). Blue points are the observations while purple lines are the model predictions.

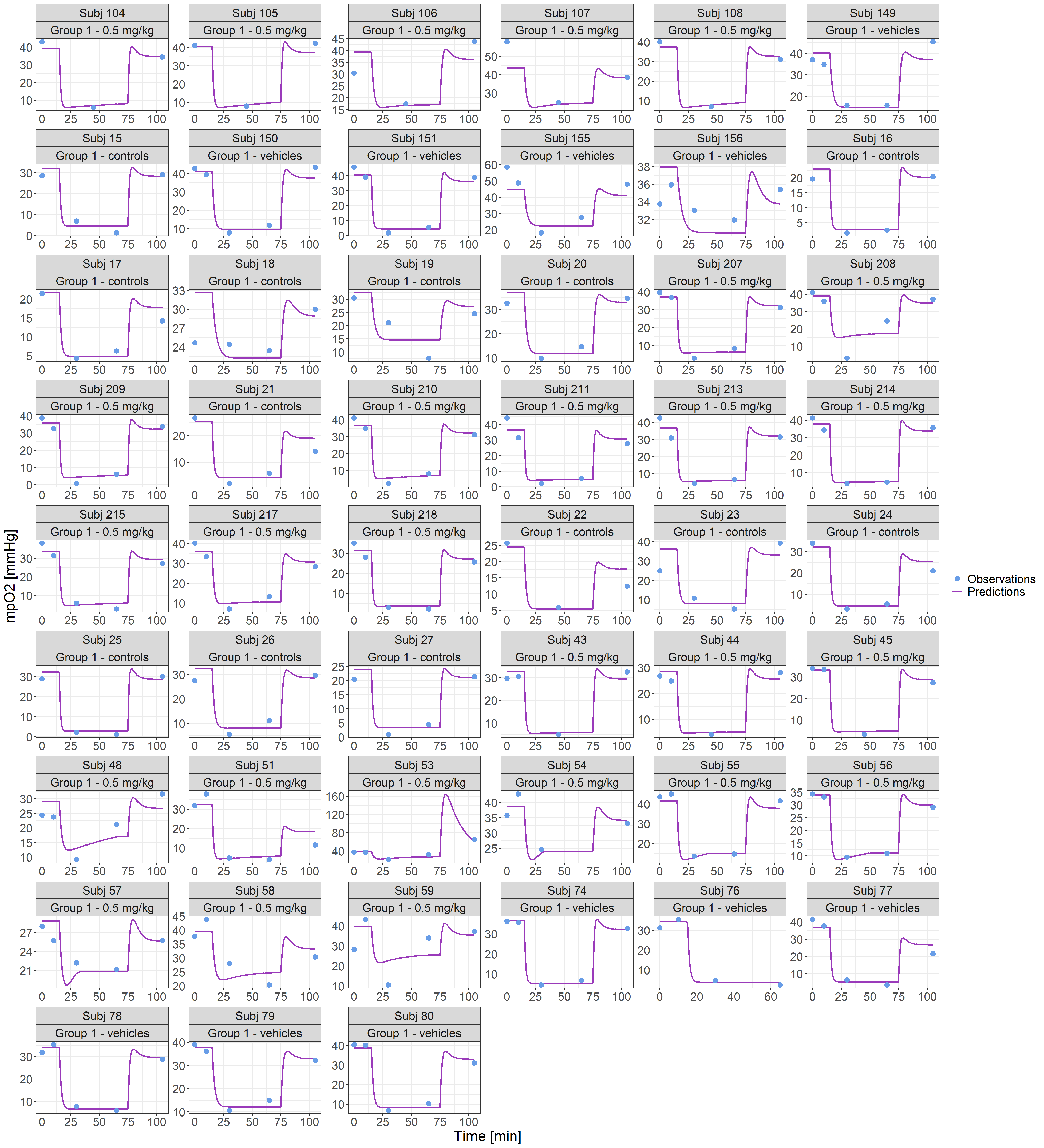

**Fig. S10** Model individual fittings of PD data in individuals belonging to experimental group 2 (drug administration 15 minutes after occlusion). Blue points are the observations while purple lines are the model predictions.

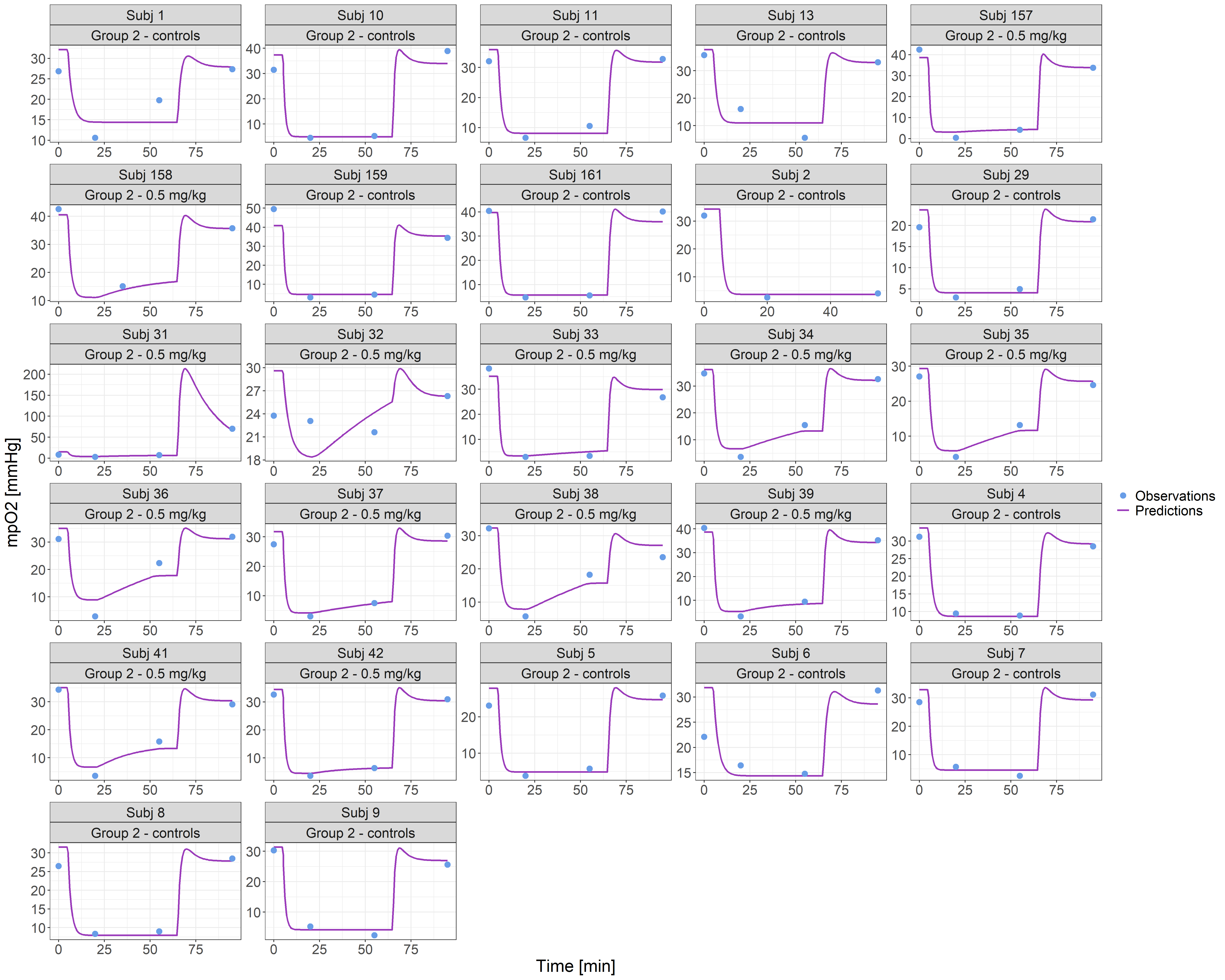

**Fig. S11** Model individual fittings of PD data in individuals belonging to experimental group 3 (drug administration 5 minutes before reperfusion). Blue points are the observations while purple lines are the model predictions.

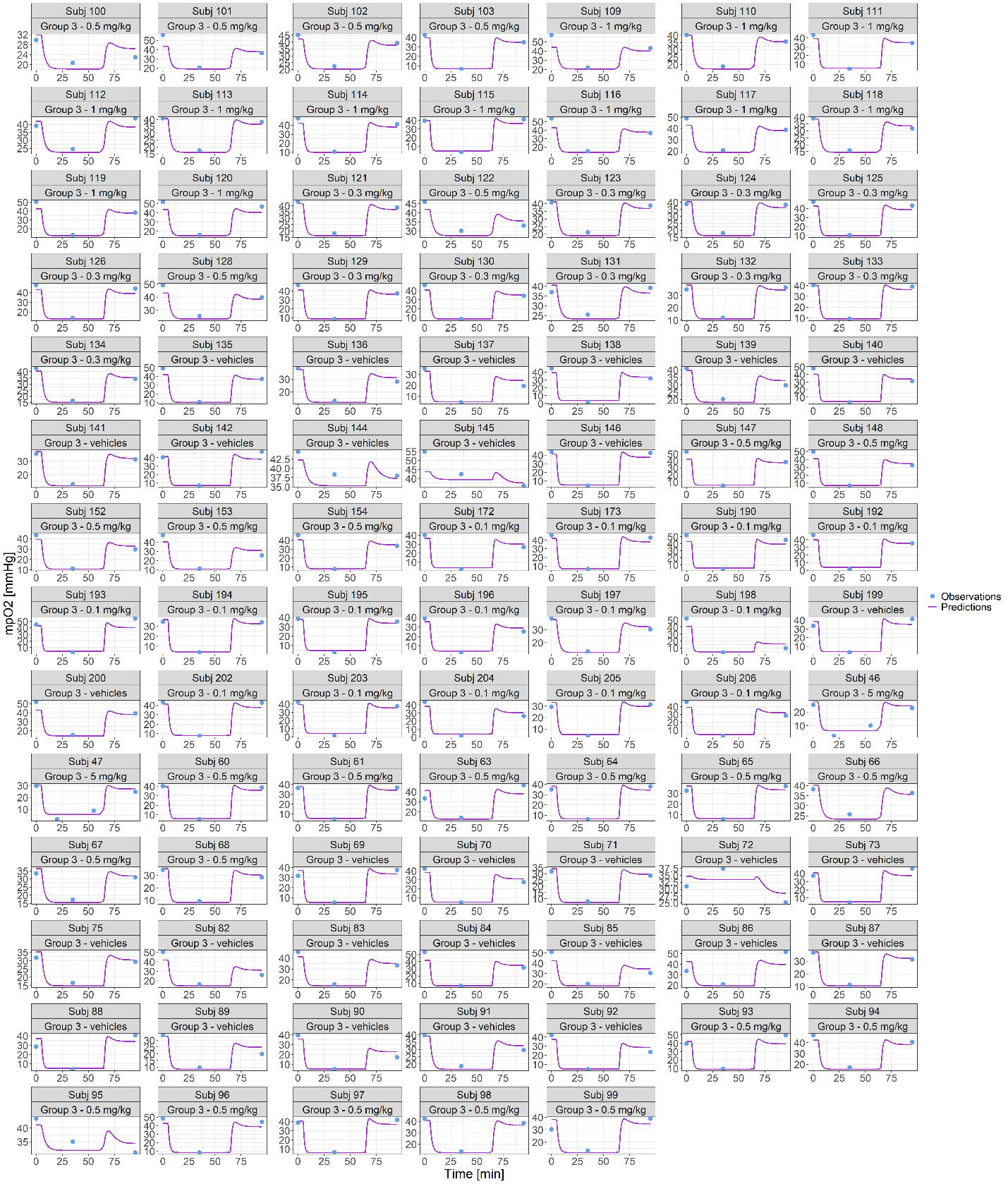

**Fig. S12** Model Goodness Of Fit plot of PD data (in Log scale). Population predictions are shown in the left box, while individual predictions are reported in the right box.

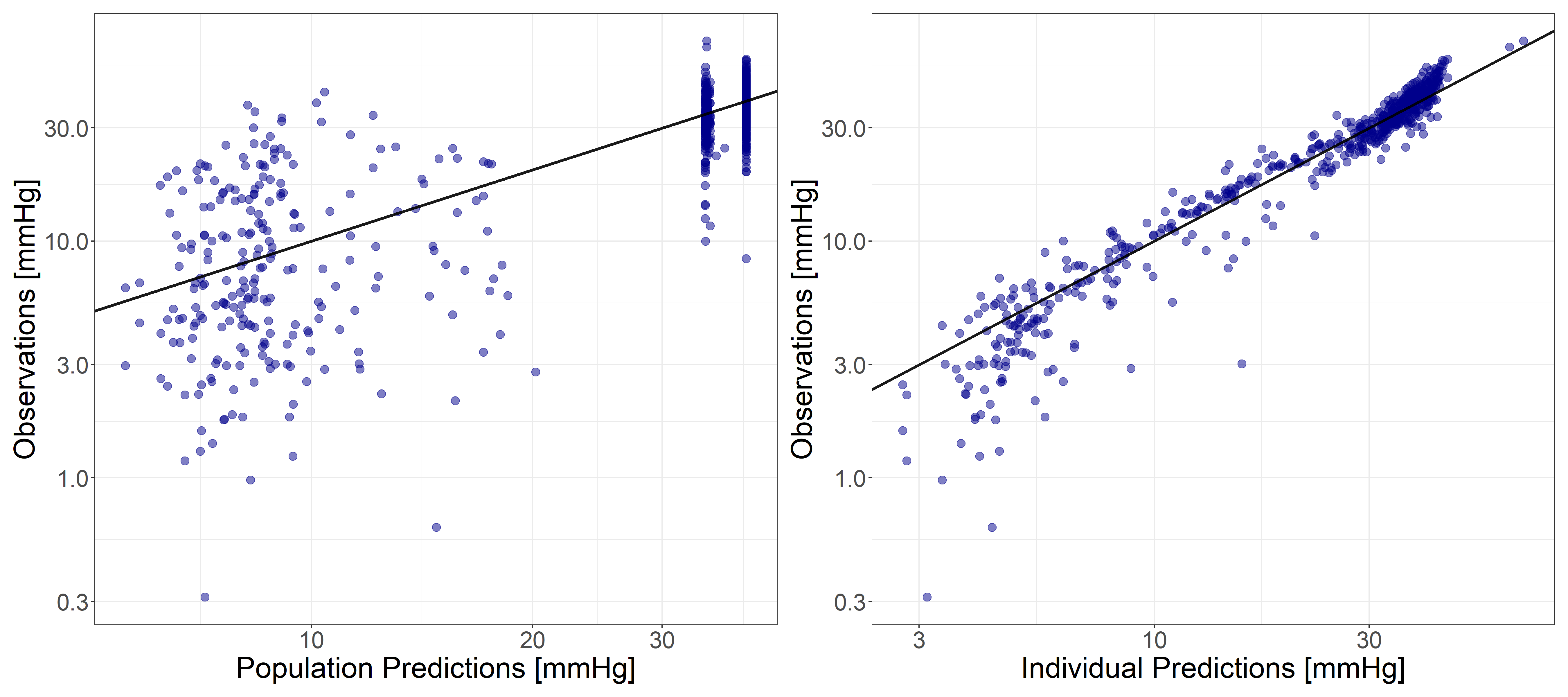
